# Exome-wide benchmark of difficult-to-sequence regions using short-read next-generation DNA sequencing

**DOI:** 10.1101/2022.11.20.517268

**Authors:** Atsushi Hijikata, Mikita Suyama, Shingo Kikugawa, Ryo Matoba, Takuya Naruto, Yumi Enomoto, Kenji Kurosawa, Naoki Harada, Kumiko Yanagi, Tadashi Kaname, Keisuke Miyako, Masaki Takazawa, Hideo Sasai, Junichi Hosokawa, Sakae Itoga, Tomomi Yamaguchi, Tomoki Kosho, Keiko Matsubara, Yoko Kuroki, Maki Fukami, Kaori Adachi, Eiji Nanba, Naomi Tsuchida, Yuri Uchiyama, Naomichi Matsumoto, Kunihiro Nishimura, Osamu Ohara

## Abstract

Next-generation DNA sequencing (NGS) in short-read mode has been recently used for genetic testing in various clinical settings. NGS data accuracy is crucial in clinical settings, and several reports regarding quality control of NGS data, focusing mostly on establishing NGS sequence read accuracy, have been published thus far. Variant calling is another critical source of NGS errors that remains mostly unexplored despite its established significance. In this study, we used a machine-learning-based method to establish an exome-wide benchmark of difficult-to-sequence regions using 10 genome sequence features on the basis of real-world NGS data accumulated in The Genome Aggregation Database (gnomAD) of the human reference genome sequence (GRCh38/hg38). We used the obtained metrics, designated “UNMET score,” along with other lines of structural information of the human genome to identify difficult-to-sequence genomic regions using conventional NGS. Thus, the UNMET score could provide appropriate caveats to address potential sequential errors in protein-coding exons of the human reference genome sequence GRCh38/hg38 in clinical sequencing.

## Introduction

Nearly 20 years after its advent, massive parallel DNA sequencing, conventionally called “next-generation sequencing (NGS),” has become a practical tool for analyzing large volumes of DNA sequences at a reasonable cost, speed, and accuracy, and is being adopted for use in clinical diagnosis. However, using NGS for diagnostic purposes requires a benchmark to validate the accuracy of NGS-based genetic testing results. Several studies have tried to establish quality control methods to maintain high-quality NGS data and benchmarking methods for internal and external precision management of NGS systems [1–5].

Variant calling using NGS is performed by mapping the obtained NGS reads onto the human reference genome, i.e., a “re-sequencing” approach. Base calling and mapping are two different sources of errors in variant calling. The Phred score is widely regarded as the gold-standard measure of base calling accuracy, with which low-quality NGS reads are filtered out to maintain a high-quality of NGS reads. In contrast, mapping accuracy is quantitatively estimated with a mapping quality score. Nonetheless, the mapping quality score depends on both the local human reference genome sequence used to map an NGS read of interest (usually 100-to 150-nt long in the case of short-read NGS) as well as the quality of the NGS read, making it difficult to directly link the mapping quality score to variant calling error rate. The mapping quality score is affected by the presence of low-mappability regions, tandem repeats, homopolymer, and other low-complexity regions (LCRs); nevertheless, quantitative measures to assess their contribution in variant calling errors in short-read NGS data are currently not established.

Recent releases of large amounts of genome-wide short-read NGS data have enabled the evaluation of variant-callingerror distribution in the human genome by short-read NGS using sequencing-by-synthesis technology [6]. The Genome Aggregation Database (gnomAD) v3.1 is a representative database of this kind, with data of 76,156 human genomes from unrelated individuals mapped against the human genome reference sequence by multiple sequencing centers using short-read NGS. The raw data from multiple sequencing centers have been reprocessed to increase the consistency of the variant calling results across sequencing centers in the gnomAD dataset. The filter information of variants in the gnomAD dataset enables the illustration of the landscape of error-proneness of current gold-standard variant-calling methods in the human genome by short-read NGS at the nucleotide-residue resolution, which revealed a distribution of difficult-to-sequence regions by short-read NGS on the basis of experimental data.

In this study, we generated a novel metric, termed the UNMET score, which allows the estimation of error-proneness of each nucleotide residue in the human genome using machine learning of the filtered information of variant data in gnomAD with 10 genomic sequence features. The UNMET score enables the identification of genomic regions that have a high possibility of sequence errors. Together with the accumulated information on structural changes in the human genome, the UNMET score would enable the accurate estimation of analytical validity of genetic testing by short-read NGS. From a practical viewpoint, the UNMET score would significantly contribute in the reporting of appropriate caveats regarding sequencing accuracy, which is recommended by the ACMG technical standard, 2021 revision [7].

## Results

### Derivation of the UNMET score

The accuracy of variant calling in high-quality NGS reads is determined by their mapping accuracy onto the reference genome sequence. Genome mappability, a measure of the similarity of regions of interest in the genome, is one of the most critical factors affecting the accuracy of variant identification by short-read NGS, but its degree of correlation with NGS accuracy is not clear [8,9]. Thus, a reductionistic approach to define difficult-to-sequence regions would be inappropriate. Instead, we tried to specify difficult-to-sequence regions using an inductive approach based on experimentally accumulated NGS data as described below.

We used the gnomAD variant dataset, which consists of genome sequencing data from more than 76,000 people with more than 6 million single nucleotide variants (SNVs) in the coding regions with allele frequency and variant quality values. We first analyzed the SNV data and its quality information in gnomAD version 3.1. To simplify the analysis, we adopted the FILTER flag (to pass or filter out data according to the quality check rule of gnomAD). As the first step, we focused only on protein-coding sequences (CDS), which resulted in 34,313,995 base positions in the GRCh38 human genome annotation for further analyses (Fig. 1A). Of these positions, 6,335,080 sites (18.5%) were identified to have at least one SNV regardless of the pass/filtered flags. Among the sites with variants, approximately 94% consisted of only passed variants while 5.3% contained filtered variants, and 0.7% contained recurrent filtered variants. Subsequently, we segregated the filtered variants in unreliable sites from the other sites. We introduced two measurements for each site: variant density (VD) and variant filter rate (VFR), depicted in Fig. 1B (described in Methods section). The two measurement metrices provide information on how many variants were observed and filtered out around a given variant site. The distribution of the VD and VFR is shown in Fig. 1C. The mean values of VD and VFR were 0.194 and 0.013, respectively, indicating that 0.194 variants were observed per site, among which 1.3% were filtered out on average. Approximately 83% of the variant positions had a VFR of 0, with no filtered variants within both 12 bp-in-length flanking regions, indicating that the position could identify variants with a high reliability. Conversely, 56,639 variant sites in CDS regions (0.8% of the variant sites) had a VFR of 1.0, indicating that all variants around these sites were filtered out, making the variants in these regions difficult to identify by short-read NGS alone. To obtain a benchmark reflecting the accuracy of variant identification using short-read NGS for each position of all CDSs at single-nucleotide resolution, we employed a machine learning approach to generate a metric designated “<u>UN</u>ified <u>MET</u>rics for unmappable, undetectable, and unreliable genomic loci” (UNMET) score for short-read NGS.

**Figure 1.**
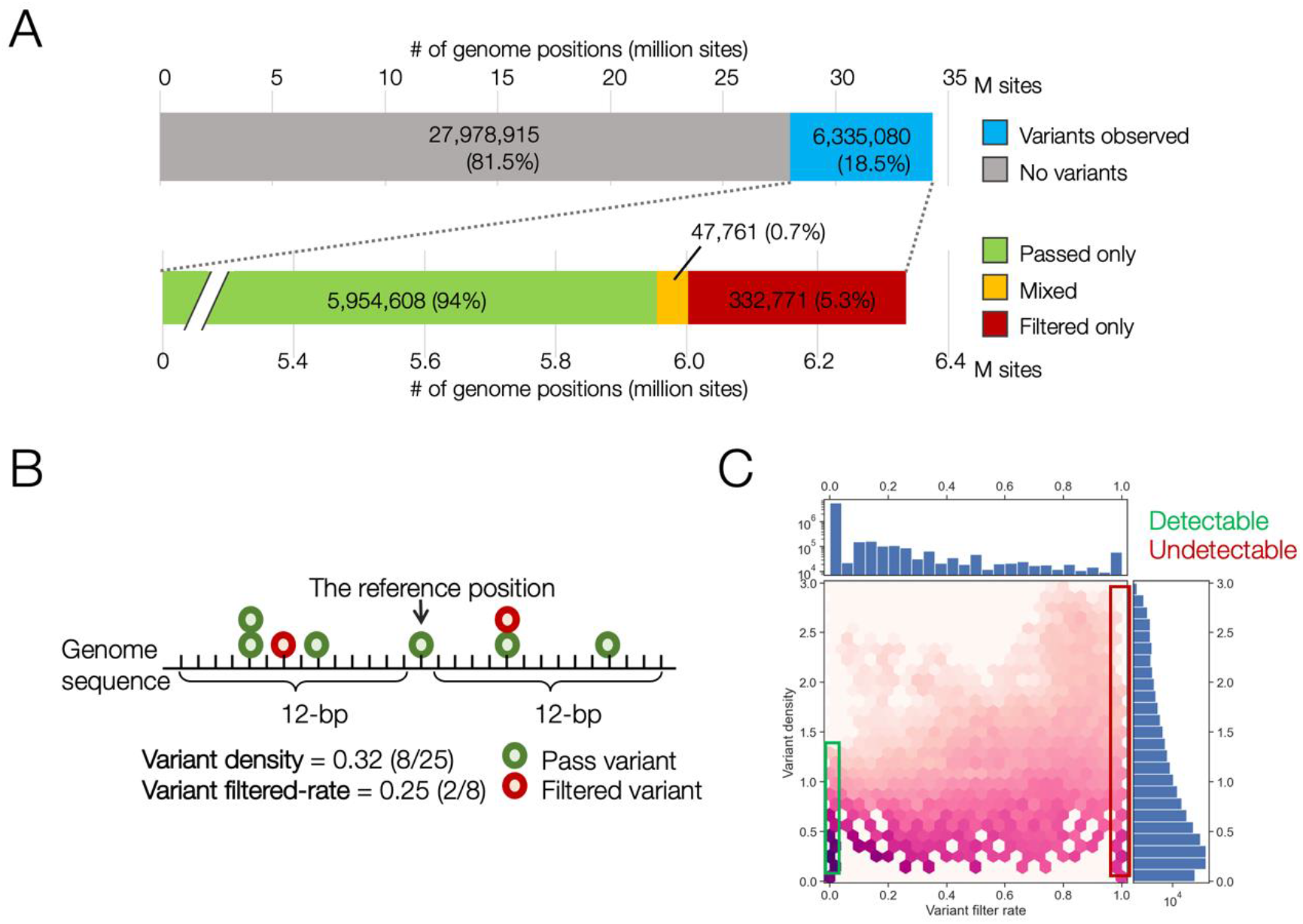
Statistics of single nucleotide variants in gnomAD version 3.1 in coding regions with filter information. (A) Proportion of genome positions with observed variants. (B) Breakdown of variant positions with filter information. Mixed indicates that two or more variants are in the same position; and both ‘passed’ and ‘filtered’ variants are observed. (C) Two-dimensional distributions of the variant filter rate (VFR; the horizontal axis) and variant density (VD; in the vertical axis) for all the sites.

The workflow of the derivation of the UNMET score is shown in Fig. 2. To train and test the score, we chose 51,715 variant sites with a VFR = 1.0 and VD > 0.3 as the negative set (“undetectable sites”), and the same number of randomly selected variant sites with a VFR - 0.0 and VD > 0.3 as the positive set (“detectable sites”). We randomly split the dataset into 80% for training and 20% for testing the machine learning protocol. We extracted 10 sequence-based features, such as genome mappability, homopolymer, tandem-repeat information as well as the standardized genome coverage depth in gnomAD data as the feature vectors (Fig. 2 and Supplementary Fig. S1A). We implemented a gradient boost decision tree algorithm, XGBoost [10], to train the model to discriminate the undetectable from the detectable sites. Evaluating the model with the testing set, we observed that the classifier could split the sites with high accuracy (area under the receiver operating characteristic curve [AUC] and Matthews’ correlation coefficient [MCC] of 0.995 and 0.962, respectively; Supplementary Fig. S1B). We then applied the prediction model to all CDS positions to assign the prediction scores, which were converted to UNMET scores, defined as percentile rank score, with a value close to 1.0 indicating that the base position was unfavorable for accurately identifying variants with a high confidence.

**Figure 2.**
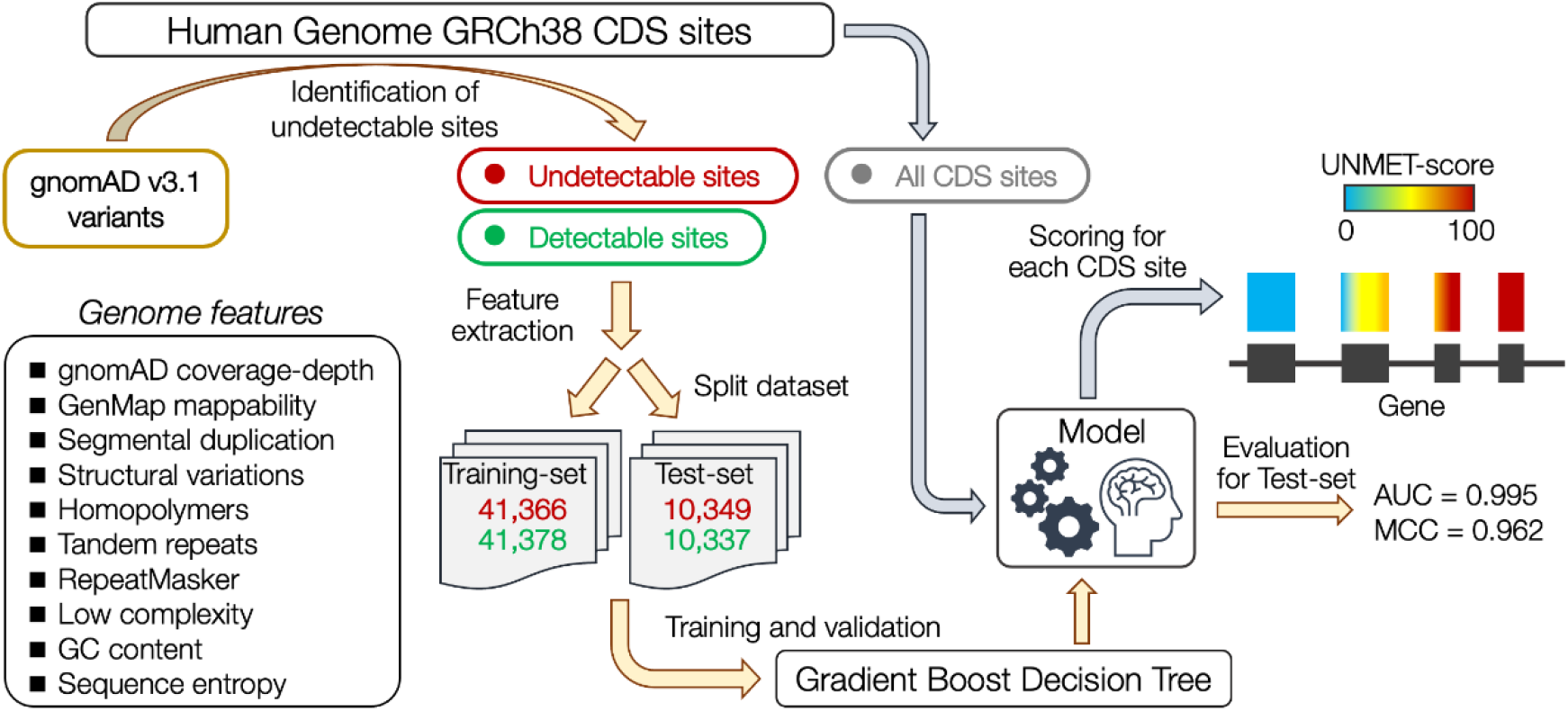
Workflow of the derivation of the UNMET score. The CDS sites on which genomic variants were located were classified into undetectable and detectable sites. Ten genomic feature vectors of the sites were used for training in machine learning to build a model that discriminates undetectable sites from detectable sites. All CDS sites were evaluated with the classifier model, and the percentile rank score of the raw score for each site was assigned as the UNMET score.

### Evaluation of the reliability of the UNMET score

#### 1. AS_VQSLOD

For evaluating the reliability of the UNMET score, we compared the UNMET scores for each genomic position to the other metrices for variant quality. First, we compared the UNMET score to the AS_VQSLOD score, a variant quality metric adopted in gnomAD v3. As expected, the UNMET score displayed an inverse correlation to the AS_VQSLOD score (Fig. 3A). The AS_VQSLOD values decreased as the UNMET score reached close to 1, and most variants with an UNMET score >0.98 were filtered out, whereas most variants at positions with an UNMET score <0.90 had passed, suggesting that the CDS positions with a high UNMET score, especially >0.97, were unreliable for genomic variant identification using short-read NGS.

**Figure 3.**
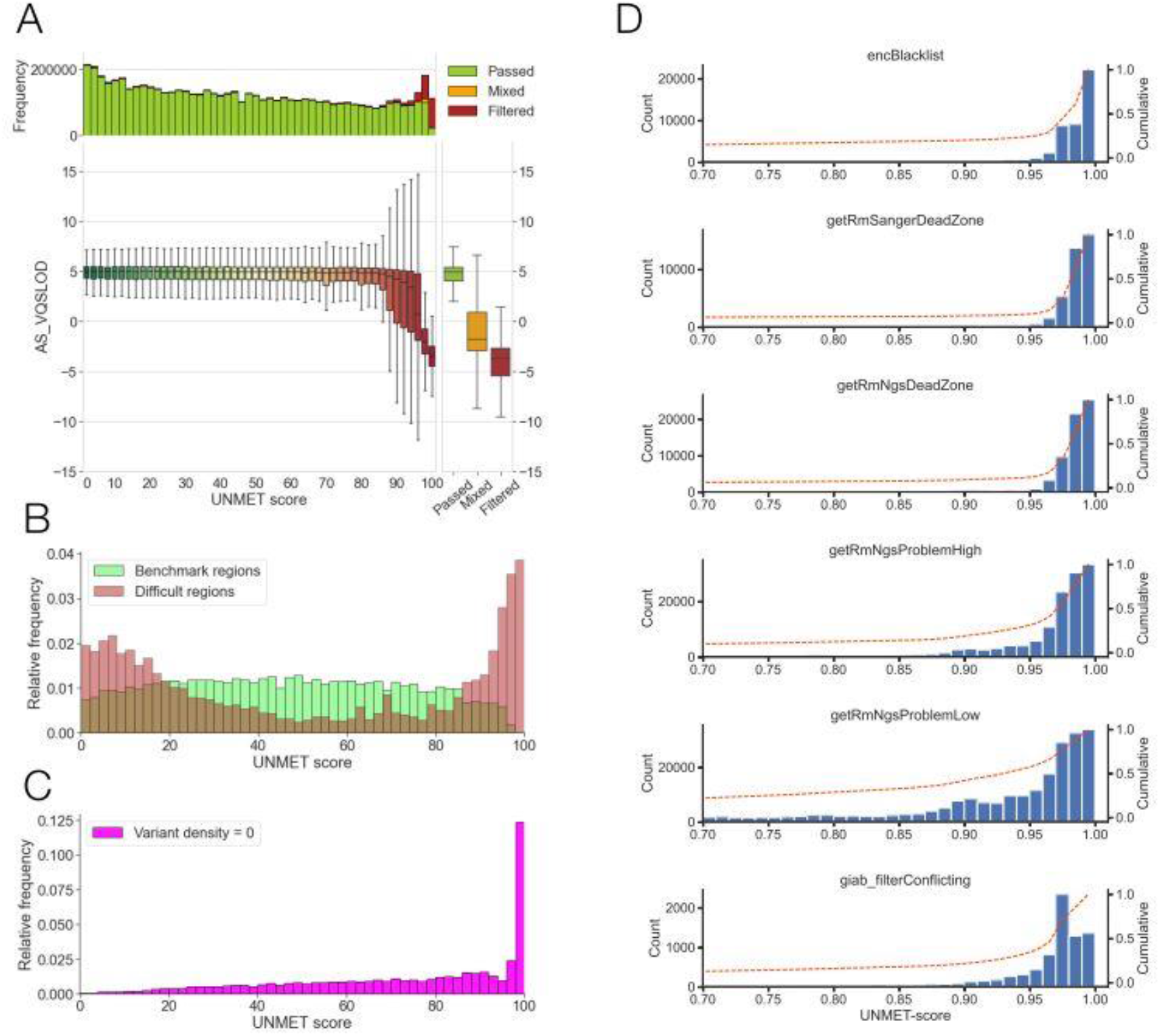
Evaluation of the reliability of the UNMET score. (A) 2D distribution plot of the UNMET score (x-axis) and AS_VQSLOD score (y-axis). The main panel depicts boxplots of AS_VQSLOD distributions in each UNMET score bin. (B) UNMET distribution in the benchmark and difficult regions designated by the GIAB dataset. (C) UNMET distribution in CDS position with variant density = 0. (D) UNMET distribution in the stratified problematic regions for NGS in the three datasets, the ENCODE Blacklist [21], GIAB [22], and NCBI GeT-RM [23], which were implemented in the UCSC Genome Browser (https://genome.ucsc.edu/).

#### 2. GIAB benchmark regions

We analyzed the distribution of UNMET scores in the CDS positions inside the benchmark and difficult-to-sequence regions in the human reference genome DNA described previously [1,2]. We compared the distribution of UNMET scores in the GIAB benchmark regions with the difficult regions where genomic variants are not reliably identified owing to technical difficulties, such as repetitive sequences (Fig. 3B). The UNMET scores in the benchmark regions were broadly distributed but most were <0.96. Notably, the score in the difficult regions showed a bimodal distribution, and one of the peaks had an UNMET score between 0.98 and 1.0, indicating that the score reasonably captured the genomic regions where genomic variants were falsely identified or misidentified with a high frequency. Moreover, approximately half of the difficult regions had an UNMET score <0.4, suggesting that these regions should be included in the benchmark regions rather than the difficult regions.

#### 3. No variant detected regions in gnomAD

Third, we analyzed the CDS positions with a VD = 0 (956,205 base positions in approximately 2.7% of the CDS positions), which denote positions with no identified SNVs in the gnomAD v3.1 dataset. There are three possible explanations for regions to have a VD = 0: first, the region could be essential for some biological functions, making its sequence relatively conserved. Second, a larger population size (>76,000) could be required to observe variants in these regions. Third, SNVs in these regions could be completely undetectable using short-read NGS. It is possible that all three situations exist in real-world data. Moreover, we observed a skewed UNMET score distribution toward the higher scores (Fig. 3C), with ∼30% of positions having an UNMET score >0.96, suggesting that variants in these regions could not be properly identified owing to technical reasons.

#### 4. Previously reported “difficult-to-sequence” regions

We analyzed the distribution of UNMET scores at nucleotide residues previously reported in the difficult-to-sequence regions (Fig. 3D). The distribution of sequence errors along UNMET scores, indicating a flexion point of error-proneness, was around 0.97. Although classification of GIAB benchmark regions (benchmark region, recurrent false positive, and VD = 0) does not illustrate a sharp discrimination of UNMET scores, it is consistent with the threshold of determined UNMET score (Fig. 3B). The genomic regions (longer than 50-nt residues) with UNMET score ≥0.97 are listed in Supplementary Table 1, which can be considered a list of “difficult-to-sequence” regions.

#### 5. Real exome data of GIAB reference genome (HG005)

We further evaluated the UNMET scores for exome sequencing of an individual sample and observed that it mimicked a practical clinical sequencing situation. We performed exome sequencing of one of the individual samples provided by the GIAB project designated as HG005 (Coriell ID, NA24631; NIST RM Number, RM8393), and the genomic variants were identified using a standard protocol assisted by the GATK v3 variant caller also used in the gnomAD dataset. We then verified the variants observed in the GIAB benchmark region with the variants confidently detected using various sequencing methodologies [2].

According to the reported variant data of HG005 [1], the grand truth variants in the benchmark region consisted of 6,915 homo- and 10,199 hetero-SNVs (Fig. 4). We compared the SNVs that we identified independently using the grand truth SNV data and labeled them as true positive (TP), false positive (FP), and false negative (FN). For homo-SNVs, most of the variants exhibited high QUAL values and were correctly detected in our exome sequencing. However, the QUAL values were slightly lower for hetero-SNVs than those for homo-SNVs, and the numbers of false-positive (293 SNVs) and false-negative variants (14 SNVs) were increased. The distribution of UNMET scores for the heterozygous SNVs was significantly different among the TP, FP, and FN variants. These results demonstrated that the UNMET score could predict the reliability of a variant correctly identified for each CDS position consistent with the real exome sequencing data.

**Figure 4.**
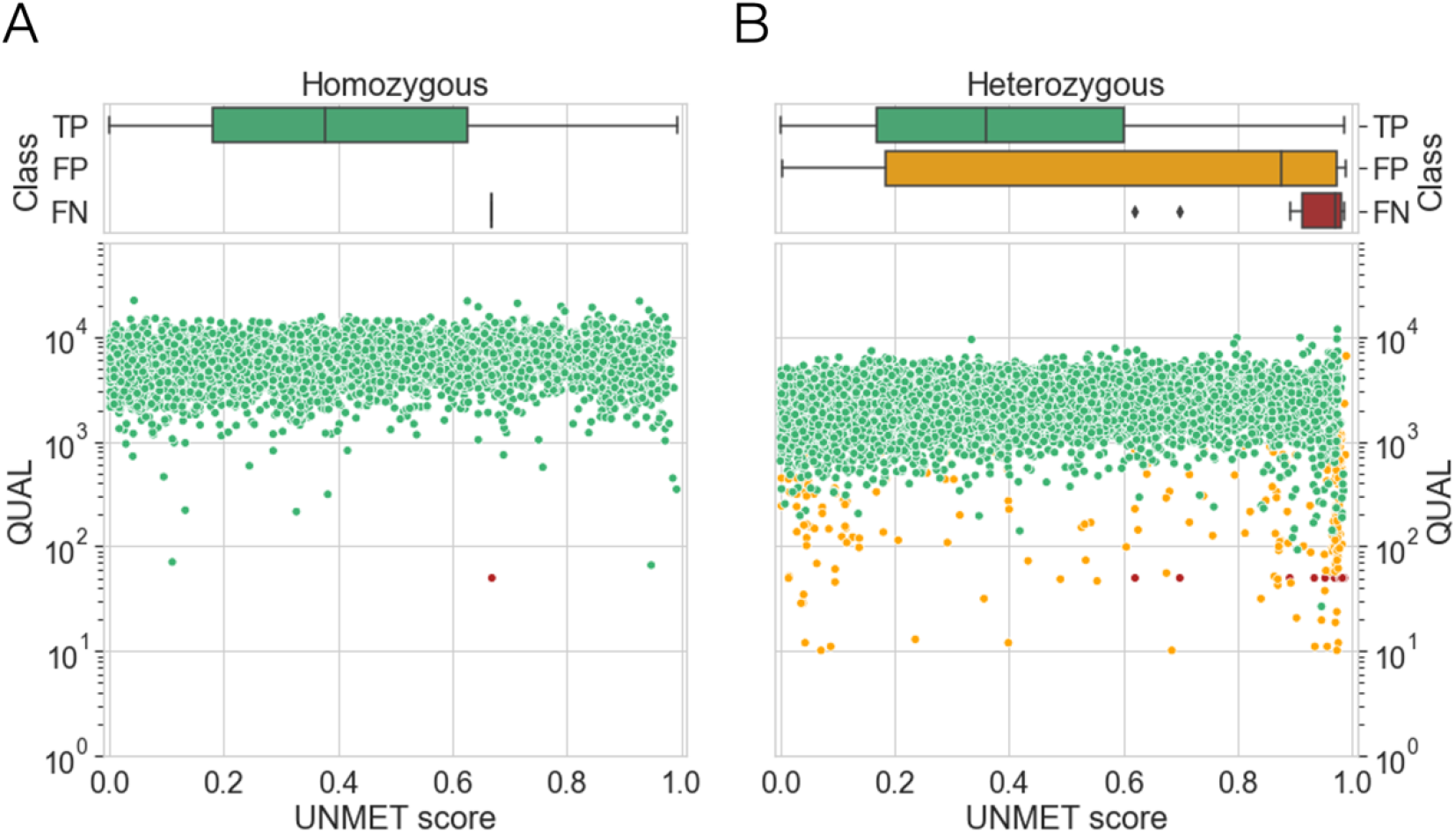
Practical evaluation of the UNMET score with exome sequencing data of the platinum genome sample. Two-dimensional distribution plot of the UNMET score (x-axis) and QUAL score (y-axis) for (A) homozygous variants and (B) heterozygous variants.

### Visualization of UNMET scores for evaluation of “error-proneness” of genomic residues in the exome regions for clinical NGS applications

The UNMET score does not directly indicate sequencing error rates like the Phred value; thus, we classified the UNMET score into “highly error-prone,” “risky,” and “reliable” regions by short-read NGS as described above. The UNMET score is a percentile rank score and thus cannot offer a clear boundary between error-prone and reliable residues. Considering this situation, we visualized the UNMET score as a heat map where highly error-prone (≥0.97), risky (0.90–0.97), and reliable (<0.90) residues are shown in red, yellow, and green, respectively, on Integrated Genomics Viewer (IGV, Fig. 5) [11,12]. For genomic regions with a high UNMET score, we also show several lines of information of genomic features together with the heat map of UNMET score: 1. gnomAD median coverage absolute z-value, 2. gnomAD exome coverage (liftover from hg19), 3. GenMap mappability (window size, 150 nucleotide long; max number of mismatches, 2), 4. Tandem repeat, 5. Homopolymeric region (>7x), 6. LCRs, and 7. GTEx exon expression levels. Additionally, we recommend visualizing actual exome data obtained by each laboratory to confirm the consistency of UNMET score with actual data under NGS conditions. For example, the exome data obtained in our NGS system (GIAB reference genome DNA, HG005) are also presented in the middle columns of Fig. 5. In the bottom part of Fig. 5, the following lines of information are given as supplements: exon–intron structure of genes, alternative loci, structural variants, and segmental duplications (classified by sequence similarity). Because the information delivered by the bottom part is useful for considering the analytical validity of genes of interest by short-read NGS, we consider it highly informative to obtain these lines of information in a single snapshot: structural variations, including recombination, duplication, and large deletion, that could cause diseases in some cases although UNMET scores derived from short-read NGS data of healthy donors do not inform about these possibilities.

**Figure 5.**
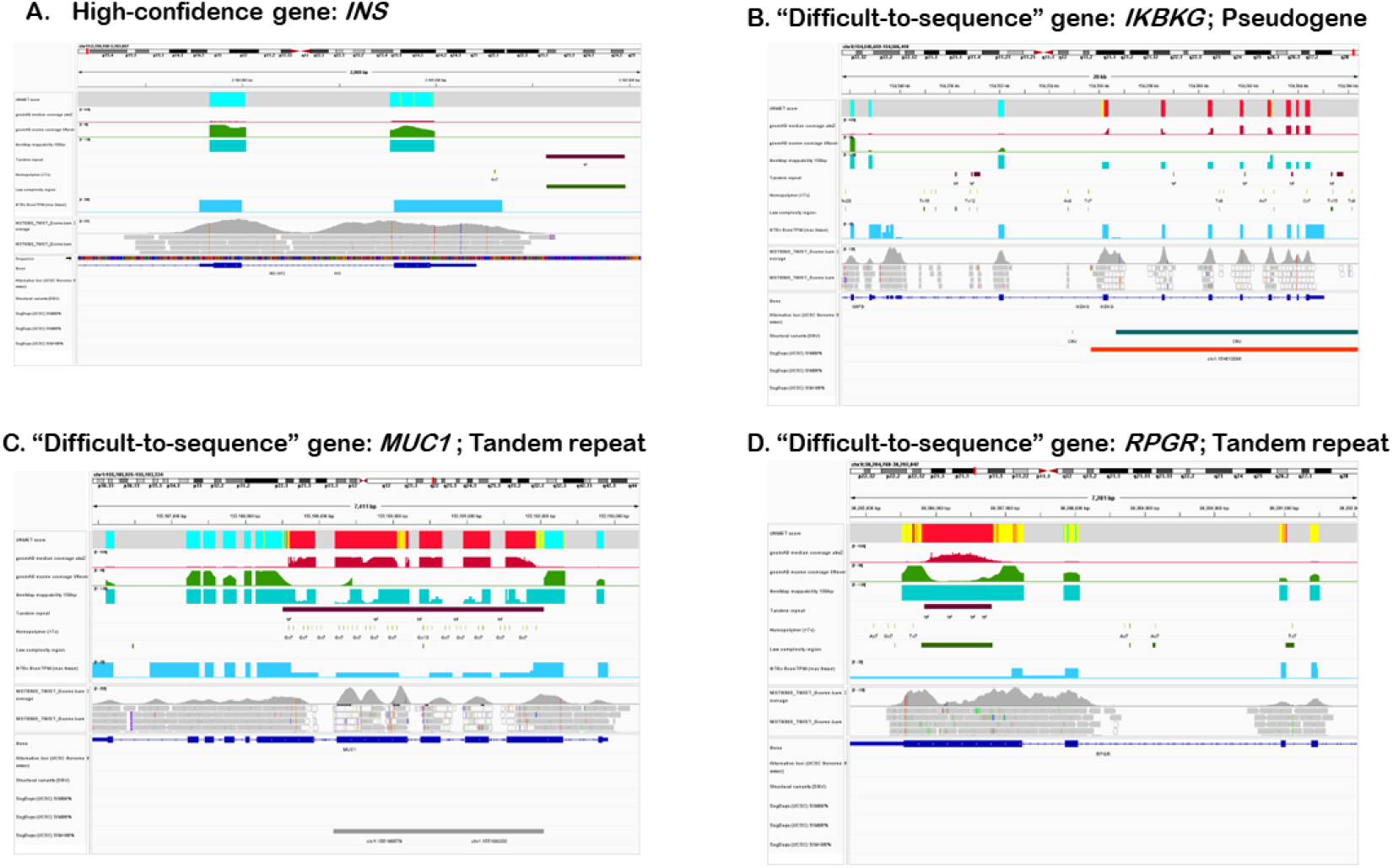
Examples showing the visualization of “difficult-to-sequence” genes on IGV. All data required for visualization on IGV as well as the session file are available from github: https://github.com/atsushihijikata/unmet-igv, aside from actual NGS exome data, which are included for confirmation of the consistency of the UNMET score with the actual NGS data under the runnning conditions employed at each laboratory. Panels exhibit examples of high-confidence gene (*INS*, panel A) and “difficult-to-sequence” genes (panel B, *IKBKG*; panel C, *MUC1*; panel D, *RPGR*).

Figure 5 demonstrates typical examples of visualization of analytical validity of genes that contain highly homologous regions to pseudogenes (Fig. 5B, for *IKBKG*), tandem repeats (Fig. 5C, for *MUC1*), or LCRs (Fig. 5D, for *RPGR*). These results indicate that this visualization tool makes it easy to overview the landscape of the error-prone regions at a glance and helps understand the cause of sequencing difficulty.

## Discussion

gnomAD is an indispensable data resource for human genetics, produced by the aggregation and harmonization of variant calls made by a well-controlled bioinformatics pipeline from a variety of large-scale sequencing projects. The variant data of gnomAD v3.1 was accumulated from 76,156 genomes from unrelated individuals, generated using multiple short-read sequencing platforms at different sequencing centers. Thus, using variant calling data from gnomAD v3.1 was suitable for the evaluation of intrinsic variant calling errors depending on local genome sequences, other than NGS technical errors, as a large number of NGS reads accumulated in gnomAD are expected to minimize the contribution of accidental NGS technical issues. Although the occurrence of NGS errors has been vaguely explained by linking several plausible causes (mapping errors and specific features of genome sequence, etc.), the gnomAD dataset enabled us to draw a solid picture of NGS error distribution by short-read NGS of the genome and sequence feature effects on NGS accuracy.

However, the UNMET score is considerably dependent on the version of the reference human genome sequence, aside from NGS running conditions (e.g., read length and DNA insert size in the library). Several variant calling discrepancies of exome sequencing exist owing to reference genome differences between GRCh37/hg19 and GRCh38/hg38 [13]. Moreover, both reference genome sequences contain false duplications even in medically relevant genes as clarified by analysis of haplotype-resolved whole genome assembly of the HG002 genome [14]. Because the training and validation data of gnomAD used in this study was based on human reference genome version GRCh38/hg38, the current UNMET score is specific for this version of the human reference genome. Thus, UNMET scores for the false-duplicated regions in each genome assembly must be re-evaluated using an appropriate genome sequence as a reference. If we accept the small uncertainty of reliability of the obtained UNMET score, it is possible for us to calculate the UNMET score from only 10 genome sequence features using machine learning data in this study. Another limitation is that we focused specifically on protein-coding exons, despite whole genome sequencing gaining popularity. An UNMET score for whole genome sequencing would require a new training dataset for machine learning because the intergenic and intronic sequence features are quite different from those in the coding regions.

Accurate clinical applications of NGS gene testing would require the identification of unexplored or undetermined regions by NGS in advance. However, because of the broad target sequences to be tested, it is difficult for clinicians and patients to estimate the missing regions by NGS in genetic testing. The UNMET score would help to identify difficult-to-sequence regions in a gene of interest and carry out additional ancillary analyses to confirm the sequences of these regions if necessary. Thus, the UNMET score established in this study would contribute to more reliable interpretation of gene testing results by NGS.

## Methods

### Definition of reliable and unreliable variant call positions and variant density in gnomAD (release 3.1)

We used the genomic variant call data in gnomAD release 3.1 to classify whether a base position was in an error-prone region and difficult to reliably call using short-read NGS equipment, i.e., an Illumina platform. The single nucleotide variant data with the filter information in all exons with its 20-bp flanking regions in protein-coding genes were extracted from the gnomAD VCF files (downloaded from the gnomAD website, https://gnomad.broadinstitute.org/downloads). To evaluate the difficulty of variant calls in genomic regions, we introduced two indicators, VD and VFR. VD was defined as the number of variants observed in a stretch *N*-bp in both ends (2*N*+1 bp window size). In the dataset, the SNVs were observed in 18.5% (6,335,080/34,313,995) of the target base positions, meaning, a variant was theoretically observed per 5 or 6 base positions. In this study, *N* was set to 12 (25-bp window size); hence, 4 or 5 variants were expected to be observed in the same window. VFR, which is the degree of filtered variants observed in a certain segment, was defined as the number of filtered variants within the same window divided by the total number of variants observed within the same region. For example, if the total number of variants observed was 8 and those of two were filtered, then the VD was 0.32 (8/25) and VFR was 0.25 (2/8), respectively. A variant position with VD ≥ 0.12 (i.e., at least three variants observed in the window) and VFR = 1.0 was considered an unreliable variant call region. In contrast, a variant position with VD ≥ 0.12 and VFR = 0.0 was considered as a reliable variant call region. The variant data in the reliable/unreliable variant regions were used for derivation of the UNMET score using machine learning.

### Sequence features for machine learning

For the machine learning approach, the genomic characteristics described below were employed as features:

1. *Genome coverage depth*. The genome coverage depth for each base position was taken from the gnomAD data and downloaded from the gnomAD website. The coverage depth of each position was normalized by the mean and standard deviation of the coverage depth in each chromosome because of the difference in the mean of the coverage depth between autosomes and non-PARs (pseudo-autosomal regions) in the × chromosome. The absolute normalized values were used for the features.
2. *Mappability*. The mappability for the human genome sequence was calculated using GenMap version 1.24 [15]. The length and mismatch parameters were set to 100 and 2, respectively. For the calculation in PAR, the sequence data of PARs in the Y-chromosome were excluded.
3. *Homopolymer tract*. The homopolymer tract (stretch of 7 or more bp) was extracted from the human genome sequences. The homopolymer tract segment with adjacent 12-bp flanking at both ends were considered.
4. *Tandem repeat*. The tandem repeat region was sought using TandemRepeatFinder (TRF) version 4.0.9 [16], and all repeat data found were treated as tandem repeats. The region with adjacent 12-bp flanking at both ends was also considered a tandem repeat region.
5. *Interspersed repeat regions*. The interspersed repeat data were obtained from the RepeatMasker data in the UCSC genome browser (https://genome.ucsc.edu/).
6. *Segmental duplication*. The segmental duplications of >1,000 bases of non-repeat masked sequences in the human genome were downloaded from the UCSC genome browser (https://genome.ucsc.edu/).
7. *LCRs*. LCRs were computed using the DustMasker program [17] implemented in the BLAST+ 2.6.0 package (ftp://ftp.ncbi.nlm.nih.gov/blast/executables/blast+). The level option was set to 30.
8. *Structural variation*. The genome regions with structural variations, including copy number variations, were obtained from the Database of Genomic Variants (DGV) [18].
9. *GC content*. The GC content 25 bp in length was calculated using an in-house Python script.
10. *Sequence entropy*. The Shannon’s information entropy for a nucleotide position was calculated based on the base frequencies in the adjacent sequences 25 bp in length.

### Training set

The unreliable and reliable genome positions defined above were considered the positive and negative set, respectively, for machine learning training. The selected positions were restricted to the CDS because of homopolymer or LCRs that are enriched in intronic regions. The number of positions for the positive and negative sets were 51,715 and 5,069,698, respectively. Because the number of positions of the negative set significantly exceeded that of the positive set, the same amount of negative data was randomly sampled from the negative set to balance the number in both sets. The dataset of 103,430 base positions with the genomic features (51,715 positive and 51,715 negative sets) were randomly split into two groups for training and testing sets in a ratio of 8:2, respectively.

### Model training

To train the model, we employed a gradient boosted decision tree algorithm (XGBoost) [10], one of the widely used machine learning algorithms, implemented in Python [19]. Hyperparameters of the model were used as a default setting. The model was trained using the training set and evaluated using the test set. Model accuracy was evaluated using AUC and MCC. The MCC is defined as:

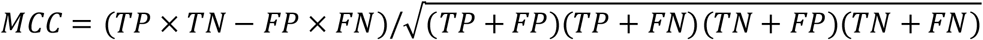

where TP, TN, FP, and FN are the ‘predicted unreliable region as unreliable,’ ‘predicted reliable region as reliable,’ ‘predicted reliable region as unreliable,’ and ‘predicted unreliable region as reliable,’ respectively.

### Test exome sequencing

Genomic DNA (NIST ID, RM8393), purchased from National Institute of Standards and Technology (Gaithersburg, MD), was used for NGS library construction using a NEXTflex Rapid DNA-Seq Kit 2.0 (PerkinElmer, Inc., Waltham, MA). Exome regions were enriched via hybridization with an IDTexome panel (Integrated DNA Technologies, Inc., Coralville, IA; The xGen Exome Research Panel v2). NGS was performed on an Illumina NextSeq2000 with a 150 nt-paired-end mode. Variants were detected using a Genome Analysis Toolkit (GATK) version 3 following the GATK Best Practices [20].

## Data availability

The UNMET score mapping data can be visualized in Integrated Genome Viewer (IGV), and the session file is freely available from github: https://github.com/atsushihijikata/unmet-igv. The exome sequencing data of human reference genome DNA (HG005) were deposited to DDBJ Sequence Read Archive (DRA, https://www.ddbj.nig.ac.jp/dra/index.html) under the run accession ID of DRR415794 (Submission Accession ID, DRA015095).

## Authors’ contributions

AH, MS, and OO drafted the manuscript. AH, MS, KK, NH, TK (Tadashi Kaname), TK (Tomoki Kosho), EN, KA, NM, and OO contributed to the conception and design of the study. SK, RM, TN, YE, YY, KM, MT, HS, JH, SI, TY, KM, YK, MF, NT, YU, and

KN validated and contributed to the improvement of visualization of the UNMET score on IGV.

## Acknowledgments

This study was motivated by a research team for a “Quality Control System for Laboratory Tests in the Field of Intractable Diseases” supported by a Health and Labor Sciences Research Grant Research Project on Intractable Diseases Policy

(JPMH19FC1018) :

