## Supplemental Figure 1 for "Exome-wide benchmark of difficult-to-sequence regions using short-read next-generation DNA sequencing"

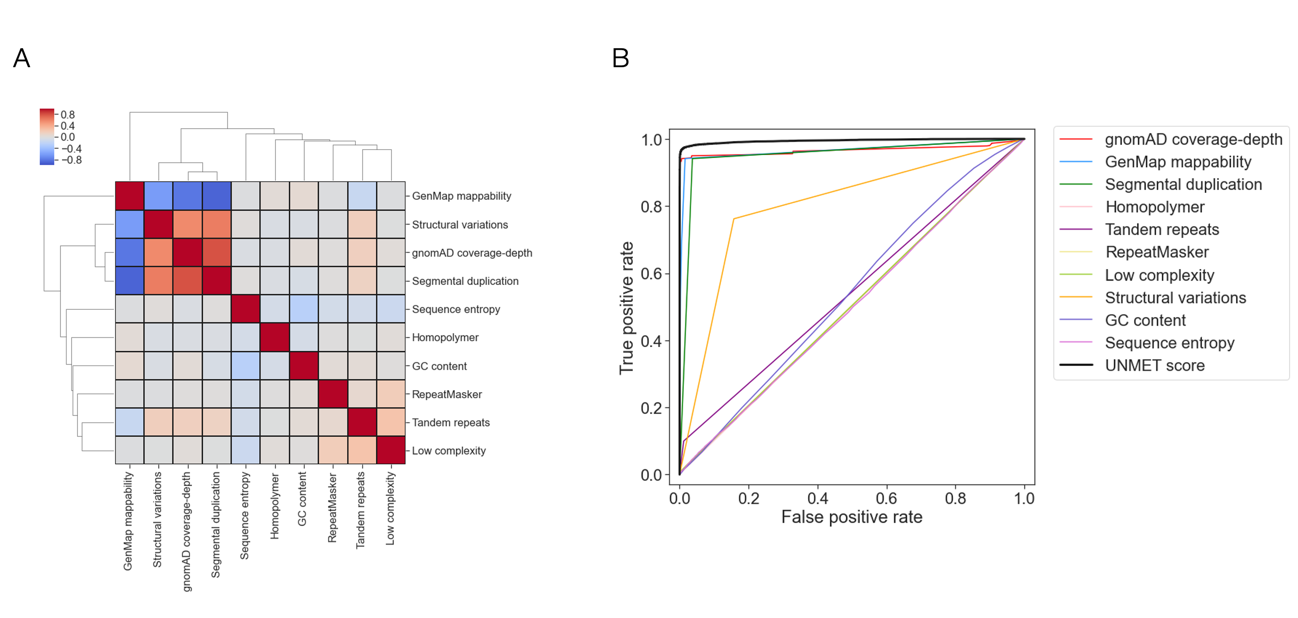

**Figure S1**. Genomic features and the use of machine learning for calculating UNMET score. (A) Heatmap of the Spearman’s correlation coefficients between the 10 genomic features used for machine learning. (B) Receiver operating characteristic (ROC) curves for evaluating the prediction accuracy. UNMET score (black) indicates the evaluation results of training with all of the features.
